# Capturing the transcription factor interactome in response to sub-lethal insecticide exposure

**DOI:** 10.1101/2020.11.26.399691

**Authors:** Victoria A Ingham, Sara Elg, Sanjay C Nagi, Frank Dondelinger

## Abstract

The increasing levels of pesticide resistance in agricultural pests and disease vectors represents a threat to both food security and global health. As insecticide resistance intensity strengthens and spreads, the likelihood of a pest encountering a sub-lethal dose of pesticide dramatically increases. Here, we apply dynamic Bayesian networks to a transcriptome time-course generated using sub-lethal pyrethroid exposure on a highly resistant *Anopheles coluzzii* population. The model accounts for circadian rhythm and ageing effects allowing high confidence identification of transcription factors with key roles in pesticide response. The associations generated by this model show high concordance with lab-based validation and identifies 44 transcription factors regulating insecticide-responsive transcripts. We identify six key regulators, with each displaying differing enrichment terms, demonstrating the complexity of pesticide response. The considerable overlap of resistance mechanisms in agricultural pests and disease vectors strongly suggests that these findings are relevant in a wide variety of pest species.

## Introduction

Insecticides are critical for control of pests in agriculture and disease vectors in public health. The intensive and widespread use of insecticides in each of these settings has led to extensive insecticide resistance (WHO, 2019), which poses a threat to both food security and global health. Vector borne diseases account for more than 17% of all infectious diseases annually (WHO, 2019), whilst around 35% of crops are lost to pre-harvest pests, underlining the importance of pesticide chemistries in global health and food security (Popp et al., 2013). Malaria control highlights the pivotal role of insecticides in global health with over 80% of the reductions in malaria cases since the turn of the century attributed to their use (Bhatt et al., 2015). Malaria control relies heavily on the distribution and use of insecticide treated bed nets (ITNs), which provide protection to the user and wider community protection through insecticide induced mortality of the adult *Anopheles* vectors (Hawley et al., 2003; Killeen et al., 2011; Killeen and Smith, 2007). All ITNs currently in use contain the pyrethroid class of insecticide; a fast-acting chemistry that induces immediate knockdown and mortality in susceptible mosquitoes. However, strength of resistance to pyrethroids is now such that populations of *Anopheles* can survive exposure with minimal effect on their life span (Hughes et al., 2020). Surviving sub-lethal exposures to pesticides is likely to have large and sustained consequences on the biology of the pest species.

Resistance to insecticides both in agricultural pests and disease vectors have been attributed to three characterised mechanisms; changes to the insecticide target site (Martinez-Torres et al., 1998; Weill et al., 2004), thickening of the cuticle to reduce penetrance (Balabanidou et al., 2016) and metabolic clearance through overexpression of detoxification protein families (Ingham et al., 2018; Müller et al., 2008; Voice et al., 2015). Recently, new resistance mechanisms have been reported (Ingham et al., 2018, 2019) and sub-lethal exposure has been shown to induce large-scale transcriptomic changes, highlighting the complexity of the insects response to insecticides (Ingham et al., 2020).

The demonstration of large-scale changes in transcriptome post-exposure emphasises the importance of transcriptional control in response to insecticide. Despite this, the induction of genes in response to insecticides is poorly studied and the regulatory processes underlying these mechanisms have remained elusive. In most important pests, cis or trans-acting regulatory elements are yet to be identified, and little published research has focused on the role of the non-coding regulatory machinery. Although recent work has identified transcription factors involved in insecticide resistance such as two transcriptional pathways: the Nrf2-cnc pathway in both disease vectors (Bottino-Rojas et al., 2018; Ingham et al., 2017) and agricultural pests (Gaddelapati et al., 2018; Kalsi and Palli, 2015) and the ARNT-AhR in agricultural pests (Hu et al., 2019; Peng et al., 2017), no studies in either setting have examined transcriptional response in a holistic manner. The availability of transcriptomic time-series data from resistant *Anopheles coluzzii* mosquitoes post-pyrethroid exposure (Ingham et al., 2020) has provided a resource to examine the importance of multiple transcription factors in response to insecticide.

Elucidating complex gene networks from transcriptomic time course data is a fundamental problem in computational systems biology (Delgado and Gómez-Vela, 2019; Jackson et al., 2020; Thompson et al., 2015). Time course data enables measurements of mRNA levels post-perturbation and allows identification of transcripts following similar expression patters over time. However, measuring changes in mRNA levels acts as a proxy for protein expression, but regulatory relationships cannot be captured by correlation alone, due to the presence of indirect regulation (gene A regulates gene B which regulates gene C), and post-transcriptional changes. To allow reconstruction of gene regulatory networks, dynamic Bayesian networks have successfully been applied to real-world time course studies (Dondelinger et al., 2013; Dondelinger and Mukherjee, 2019; Murphy and Mian, 1999), allowing identification of key regulatory pathways within a system. As circadian rhythms can play a significant role in gene expression patterns over short time-scales (Rund et al., 2011), these models additionally allow correction for sinusoidal patterns with 24-hour period.

Here, we apply a modified dynamic Bayesian network method to whole-organism microarray data taken at ten time-points post exposure to a pyrethroid insecticide. The method corrects for both circadian patterns and mosquito ageing, which have previously been shown to be important in the insecticide resistance phenotype (Jones et al., 2012; Rund et al., 2016). The Bayesian network approach allows identification of key regulatory factors influencing the expression of transcripts in response to insecticide exposure. Based on validation experiments, we estimate that the inferred network has 70% precision, indicating strong concordance of experimental data to model prediction. The network is made freely available through a ShinyR application, allowing non-bioinformaticians to easily access and visualise the data. Several transcription factors are highlighted as potential key regulators in response to pyrethroid insecticide. This study demonstrates the importance and complexity of transcriptional control of insecticide response, which is likely to have cross species applicability due to relative conservation of transcriptional pathways (Hsia and McGinnis, 2003) and near total overlap of resistance mechanisms.

## Results

### Identification of transcription factors involved in insecticide resistance

Of the transcripts in the *Anopheles* microarray, approximately 4% are putative transcription factors, based on *Drosophila* homology. As exploration of all possible transcription factor/transcript associations was not computationally feasible, the number of transcription factors had to be reduced to <50. Of the 559 total transcription factors, 44 were used in further analyses (Table 1). These transcription factors were selected based on resistance-associated GO term enrichments in transcription factor-transcript clusters (Zhang et al., 2018) found using a previously published library of microarray data comparing resistant and susceptible *Anopheles* species across Africa (Ingham et al., 2018). A number of these transcription factors have known roles in stress response in *Drosophila* (Table 1); however, only *Maf-S, Met* and *Dm* have previously been linked with insecticide response in mosquitoes (Ingham et al., 2018, 2017). Of the transcription factors selected for analysis the following have been studied in mosquitoes: *p53* has been shown to respond to arboviral infection (Chen et al., 2017); *Rbsn-5* has been shown to be involved in egg shell formation (Amenya et al., 2010); *l(1)sc* is linked with sensory tissue development (Wülbeck and Simpson, 2002); *kaya*k is involved in salivary gland response to arboviral infection through *JNK* pathway activation (Chowdhury et al., 2020); *Hnf4* is linked to ecdysone and *Met* mediated lipid metabolism (Wang et al., 2017); *Cyc* controls the circadian ryhthm (Maliti et al., 2016); *REL1* and *REL2* are involved in immune response (Luna et al., 2006); *Kr-h1* is essential for egg development (Fu et al., 2020) and *Pan* is linked with chromatin changes upon *Plasmodium* infection (Ruiz et al., 2019).

**Table 1:** List of Transcription Factors included in further analysis. 44 Transcription factors included in the implementation of the dynamic Bayesian model, including VectorBase Transcript ID, Drosophila gene name, FBgn identifier, % identity (taken from VectorBase), putative function and network interactor summary KEGG/GO enrichment from this study.

| Transcript ID | Gene Name | Homolog | % Identity | Role in Drosophila | Citation | Network Enrichment |
| --- | --- | --- | --- | --- | --- | --- |
| AGAP000057-RA | shaven (sv) | FBgn0005561 | 34.12 | Sensory tissue development | Kavaler et al. 1999 (Kavaler et al., 1999) | Hippo Signalling |
| AGAP000066-RA | Sox102F | FBgn0039938 |  | Neuronal development, behaviour and Wnt signalling | Li et al. 2017 (Li et al., 2017) | Protein Catabolism, Tissue Regeneration |
| AGAP000141-RA | CG31224 | FBgn0051224 | 17.03 | Unknown |  | Autophagy, longevity |
| AGAP000547-RA | Rbsn-5 | FBgn0261064 | 42.29 | Endosome assembly | Morrison et al 2008 (Morrison et al., 2008) | RNA-related, Transferase Activity |
| AGAP000646-RA | Diminutive (Dm, dMyc) | FBgn0262656 | 13.21 | Glucose and lipid metabolism, development | Parisi et al. 2013 (Parisi et al., 2013) | Sugar and Lipid Metabolism, Miscellaneous Metabolism |
| AGAP000876-RA | achaete-scute complex (l(1)sc) | FBgn0002561 | 26.42 | Neuronal development, dopaminergic neurons | Stagg et al 2011 (Stagg et al., 2011) | UDP-glycosyltransferase, Drug Metabolism |
| AGAP001093-RA | kayak (kay) | FBgn0001297 | 30.06 | JNK signalling, wound healing, neuronal development | Ramet et al. 2002 (Rämet et al., 2002); Miotto et al. 2006 (Miotto et al., 2006) | RNA-related Processes |
| AGAP001156-RA | PSEA-binding protein 95kD (Pbp95) | FBgn0037540 | 13.89 | Small nuclear RNA activating complex | Li et al 2004 (Li et al., 2004) | Signalling Pathways |
| AGAP001388-RA | Doublesex-Mab related 93B (dmrt93B) | FBgn0038851 | 41.61 | Mouth development | Panara et al 2019 (Panara et al., 2019) | Longevity-related |
| AGAP001786-RA | Osa | FBgn0261885 | 36.83 | EGFR signalling | Terriente-Feliz and de Celis 2009 (Terriente-Félix and de Celis, 2009) | Transcription Factors |
| AGAP001994-RA | Brahma associated protein 111kD (Bap111) | FBgn0030093 | 40.1 | Chromatin remodelling | Papoulas et al. 2001 (Papoulas et al., 2001) | Miscellaneous Metabolism |
| AGAP002082-RA | Squeeze (sqz) | FBgn0010768 | 35.47 | Neuronal development | Terriente-Feliz et al 2007 (Félix et al., 2007) | Transcription Factors |
| AGAP002155-RA | Hepatocyte nuclear factor 4 (Hnf4) | FBgn0004914 | 52.85 | Lipid mobilisation, glucose homeostasis and mitochondrial function | Palanker et al. 2009 (Palanker et al., 2009); Barry and Thummel 2016 (Barry and Thummel, 2016) | Lipid Transporters, Galactose Metabolism |
| AGAP002352-RB | p53 | FBgn0039044 | 14.2 | Genotoxic stress response | Brodsky et al. 2004 (Brodsky et al., 2004) | Glycolysis |
| AGAP002773-RA | Stripe (sr) | FBgn0003499 |  | Muscle development | Lee et al. 1995 (Lee et al., 1995) | Endosome-related |
| AGAP002902-RA | Medea (Med) | FBgn0011655 | 52.42 | Muscle development through BMP and dpp Pathways | Wisotzkey et al. 1998 (Wisotzkey et al., 1998) | Ribosome-related |
| AGAP002920-RA | CG17829 | FBgn0025635 | 17.84 | Unknown |  | Protein Complex Binding, DNA Repair |
| AGAP002954-RA | Cell division cycle 5 (Cdc5) | FBgn0035136 | 63.63 | Spliceosome | Herold et al. 2009 (Herold et al., 2009) | FoxO Signalling, Sugar- and Cytoskeletal- related Processes |
| AGAP003117-RA | Capicua (cic) | FBgn0262582 | 19.37 | EGFR, Torso and TOLL signalling | Astigarraga et al. 2007 (Astigarraga et al., 2007); Papagianni et al.2018 (Papagianni et al., 2018) | Neuronal-related Processes |
| **AGAP003449-RA** | Rootletin (Root) | FBgn0039152 | 46.08 | Hearing, touch and taste | Chen et al. 2015 (Chen et al., 2015) | Steroid Biosynthesis, Peroxisome |
| **AGAP003669-RA** | Drop (Dr) | FBgn0000492 | 61.4 | Eye and nerve development | Tearle et al. 1994 (Tearle et al., 1994) | Endopeptidase-related |
| **AGAP004864-RA** | Protein on ecdysone puffs (Pep) | FBgn0004401 | 38.87 | Hsp70 response through hnRNP complex | Hamann et al. 1998 (Hamann and Strätling, 1998) | None |
| **AGAP004990-RA** | Multiprotein bridging factor 1 (mbf1) | FBgn0262732 | 74.15 | Co-activator to induce stress-response genes | Jindra et al. 2004 (Jindra et al., 2004) | Translation-related Processes |
| **AGAP005437-RA** | Inverted repeat binding protein 18 kDa (Irbp18) | FBgn0036126 |  | Inhibitor of the conserved stress response protein dATF4/Crc | Blanco et al 2020 (Blanco et al., 2020) | Longevity |
| **AGAP005551-RA** | Rabaptin-5-associated exchange factor for Rab5 (Rabex-5) | FBgn0262937 | 37.75 | Ras pathway homeostasis | Yan et al. 2010 (Yan et al., 2010) | ABC-transporters |
| **AGAP005641-RA** | CG9705 | FBgn0036661 | 54.78 | Sensory neurons | Iyer et al. 2013 (Iyer et al., 2013) | Protein Sorting |
| **AGAP005655-RA** | Cylce (Cyc) | FBgn0023094 | 35.25 | Circadian rhythm | Rutila et al. 1998 (Rutila et al., 1998) | Peptidases, Drug Metabolism, Digestion |
| **AGAP006022-RA** | Methoprene tolerant (Met) | FBgn0002723 | 21.2 | Juvenile hormone binding | Jindra et al. 2015 (Jindra et al., 2015) | Peroxisome |
| **AGAP006061-RA** | Ken | FBgn0000286 | 5.92 | JAK/STAT pathway | Arbouzova et al. 2006 (Arbouzova et al., 2006) | Proteolysis |
| **AGAP006392-RA** | CG4617 | FBgn0029936 | 38.58 | Unknown |  | Autophagy |
| **AGAP006601-RA** | MEP-1 | FBgn0035357 | 31.69 | Chromatin remodelling | Reddy et al. 2010 (Reddy et al., 2010) | MAPK-Signalling |
| **AGAP006642-RA** | Defective proventriculus (dve) | FBgn0020307 | 47.98 | Mitochondrial reactive oxygen species modulator | Baqri et al. 2014 (Baqri et al., 2014) | Mitochondria-related |
| **AGAP006736-RA** | Sugarbabe (sug) | FBgn0033782 | 28.24 | Regulation of lipid and carbohydrate metabolism | Varghese et al. 2010 (Varghese et al., 2010) | Lipid-related Processes |
| **AGAP006747-RA** | Relish (REL2) | FBgn0014018 | 24.12 | Immune response | Dushay et al. 1996 (Dushay et al., 1996) | None |
| **AGAP009444-RA** | Suppressor of variegation 205 (Su(var)205) | FBgn0003607 | 23.47 | Hsp70 response through activation of euchromatic genes | Piacentini et al. 2003 (Piacentini et al., 2003) | Valine, Leucine and Isoleucine Degradation |
| **AGAP009494-RA** | Ets at 21C (Ets21C) | FBgn0005660 | 34.25 | Stress inducible transcription factor through JNK | Mundorf et al. 2019 (Mundorf et al., 2019) | Glutathione, Oxidative Phosphorylation |
| **AGAP009515-RA** | REL1 | FBgn0260632 | 38.96 | Toll pathway | Gross et al. 1999 (Gross et al., 1999) | Miscellaneous Metabolism |
| **AGAP009662-RA** | Kruppel Homolog 1 (Kr-h1) | FBgn0028420 | 36.47 | 20-hydroxyecdysone linked | Pecasse et al. 2000 (Pecasse et al., 2000) | Drug Metabolism |
| **AGAP009676-RA** | Chameau (chm) | FBgn0028387 | 34.66 | JNK signalling | Miotto et al. 2006 (Miotto et al., 2006) | DNA Repair |
| **AGAP009888-RA** | CG33695 | FBgn0052831 | 53.3 | Unknown |  | Degradation-related |
| **AGAP009899-RA** | klumpfuss (klu) | FBgn0013469 | 42.86 | Cell death, mitochondrial function, EGFR signalling | Protzer etl al. 2008 (Protzer et al., 2008); Chen et al. 2008 (Chen et al., 2008) | Respiratory-related, Mitochondria-related, Drug Metabolism |
| **AGAP009983-RA** | Net | FBgn0002931 | 35.88 | EGFR signalling | Terriente-Feliz and de Celis 2009 (Terriente-Félix and de Celis, 2009) | Terpenoid Biosynthesis |
| **AGAP010405-RA** | Maf-S | FBgn0034534 | 63.7 | Reactive oxygen species stress response | Misra et al. 2011 (Misra et al., 2011) | Mitophagy |
| **AGAP012389-RA** | Pangolin (Pan) | FBgn0085432 | 24.47 | Wingless signalling | Brunner et al. 1997 (Brunner et al., 1997) | Peptidase-activity, Chromatin-related |

### Modelling the insecticide response network

To explore the role of the identified transcription factors in insecticide response, a previously generated time course experiment comparing pyrethroid exposed and unexposed *Anopheles coluzzii* was used (Ingham et al., 2020). In this dataset 8832 transcripts (9316 probes) were significantly differentially expressed in at least one of the ten time points (from immediately post-exposure to 72 hours post-exposure), demonstrating large changes to mosquito biology post-exposure. This dataset was then used to model the gene regulatory relationships using a dynamic Bayesian network (DBN) approach (Dondelinger et al., 2013) which infers the regulators of each transcript from the set of selected transcription factors using the time-course of log-fold changes compared to the unexposed baseline measurement, correcting for ageing and circadian rhythms. A Markov chain Monte Carlo (MCMC) algorithm was used to draw samples from the posterior distribution of the network model given the data, and then associations were ranked between target genes and transcription factors using the marginal posterior probability of the corresponding edge (defined as a predicted transcription factor – transcript association) in the network. Since experimental validation of all discovered edges is prohibitively expensive, an important consideration was how many associations needed to be tested in order to establish the validity of the network inference approach. A simulation study was performed under the assumption that the number of genes regulated by each transcription factor follows a Poisson distribution with mean 10. We showed that under some assumptions (see Materials and Methods) testing 4 regulatory relationships for each of 7 transcription factors has a 70% chance of obtaining an estimate of the precision that falls within 10% of the true precision, and a 95% chance of obtaining an estimate that falls within 20% of the true precision. For 5 transcription factors with 4 regulatory relationships, this still gives a 65% chance of an estimate within 10%, and a 90% chance of an estimate within 20% of the true precision.

The model was validated using quantitative PCR to confirm the interactions predicted by the model. Successful dsRNA mediated knock down was performed on 5 transcription factors, these showed knock down 48-hours post insecticide exposure (Supplementary Figure 1); the single time point used for model confirmation (Figure 2, Supplementary Table 2). Four transcript interactors were chosen randomly for each transcription factor based on a posterior probability of > 0.2. To determine the change in transcript expression post-exposure and to determine whether predicted interactors were influenced by the knock down of the stated transcription factor 2 comparisons were made: (i) GFP-injected exposed vs GFP-injected unexposed and (ii) Exposed transcription factor knockdown compared to exposed GFP-injected for the two comparisons respectively (Supplementary Table 2). Of the 20 interactors (5 transcription factors × 4 interactors), 14 demonstrated concordance with the model, showing a substantial change in expression due to transcription factor knockdown, indicating 70% model precision (Figure 2, Supplementary Table 2).

**Figure 1:**
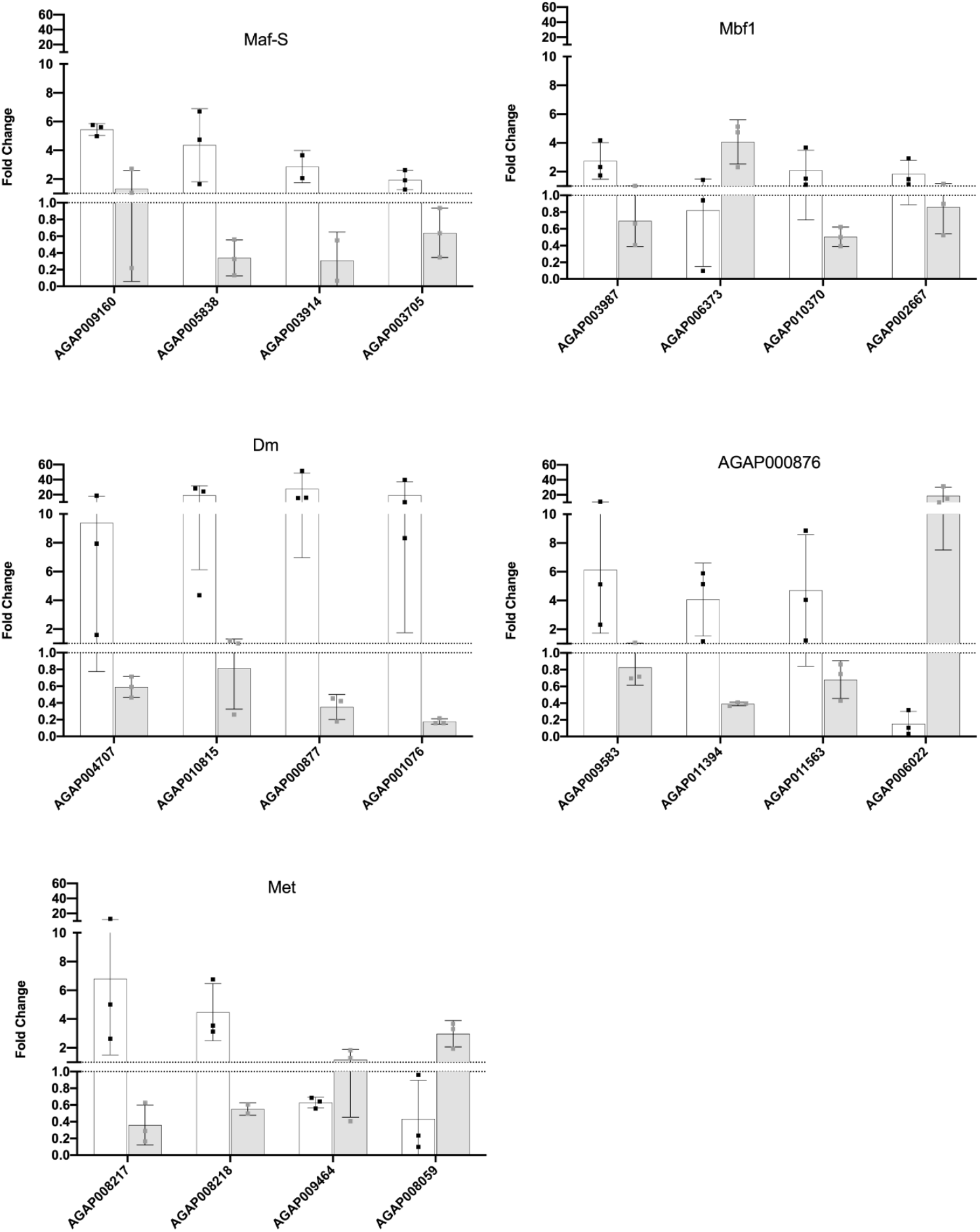
Model validation. mRNA fold change (y-axis) of each transcript (x-axis) for each transcription factor showing knockdown 48-hours post-deltamethrin exposure. White bars show qPCR results from GFP-injected exposed mosquitoes (48-hours post exposure) compared to GFP-injected unexposed mosquitoes (48-hours post injection) to show induction effect in absence of treatment and grey bars show transcription factor-injected exposed (48 hours post exposure) vs GFP-injected exposed mosquitoes (48-hours) to demonstrate the effect of transcription factor knockdown. Error bars show standard deviation.

**Figure 2:**
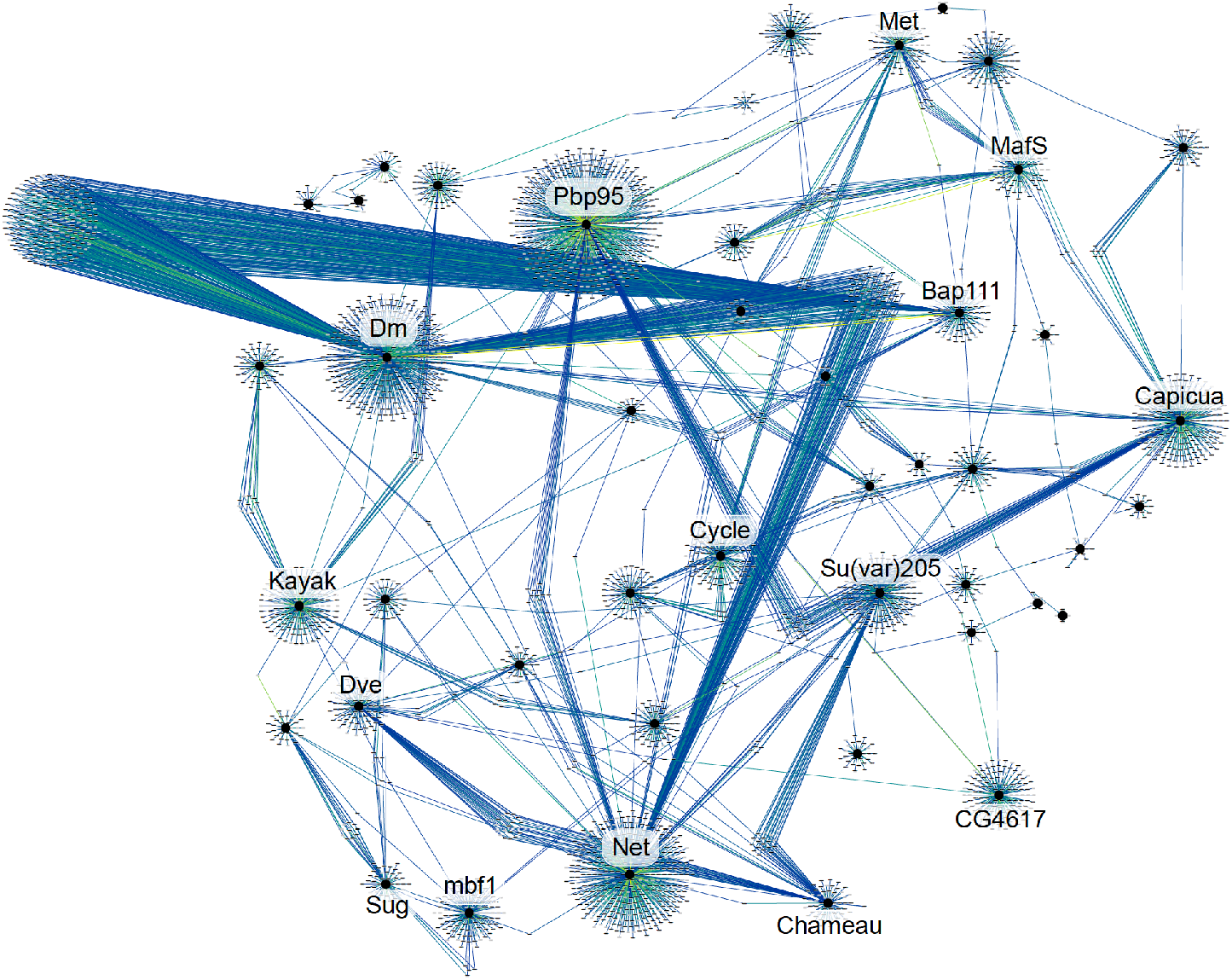
Network overview. Emboldened black circles represent all 44 transcription factors, with grey nodes representing associated transcripts. Directed edges are coloured on posterior probability gradient from dark blue (0.25) through green (0.5) to yellow (0.75). High posterior probability indicates higher confidence in the interaction. The 14 hub transcription factors, with > 100 associations are labelled.

### Network Overview

In order to determine what the optimal cut-off for the marginal posterior probability values should be, a permutation test was performed whereby the observed log-fold values for one of the 44 transcription factors are randomly permuted, so that the original time associations were no longer present. Any association between this transcription factor and the target gene would then be purely due to chance. This process was then repeated 500 times, inferring the edges for all 44 transcription factors each time. The resulting marginal posterior probability values were then analysed for the randomised transcription factor and showed that a threshold of 0.23 was sufficient to only produce one false positive out of 500 randomizations, or a false positive rate of 0.002. To be conservative, we used a cut-off of 0.25 which resulted in assignment of 4132 transcripts to the 44 transcription factors.

The complete network using a posterior probability cut-off of 0.25 is displayed in Figure 1. Due to the constraints imposed by this model on number of parent nodes tested, simple network descriptive data was generated only for edges from the selected transcription factors. The average edge count was 107.84±150.42 demonstrating high variance in connectivity as seen in Figure 1 with a range of 5 associations to 702. Fourteen transcription factors are network hubs, defined as nodes with a high number of associations (>100) (Table 2), including *Dm, Met* and *Maf-S* all previously linked with the insecticide resistance phenotype (Ingham et al., 2018, 2017) and *mbf1* a stress response transcription factor (Jindra et al., 2004). To enable the network to be freely accessible an application NetworkVis has been written in ShinyR (Chang et al., 2017) and is freely available (https://github.com/VictoriaIngham/NetworkVis_TimeCourse) with all associated data. Users can manually select a posterior probability cut-off between 0.1-0.8, select and rearrange nodes and edges in the network and identify *a priori* transcription factors through visual means rather than working with a large text file.

**Table 2:** Transcription factor hubs. Identifier, gene name and number of associations for 14 transcription factor hubs within the network.

| Transcription Factor | Drosophila Homolog | Number of associations |
| --- | --- | --- |
| AGAP000646-RA | Diminutive | 702 |
| AGAP001156-RA | Pbp95 | 591 |
| AGAP009983-RA | Net | 490 |
| AGAP001994-RA | Bap111 | 323 |
| AGAP003117-RA | Capicua | 269 |
| AGAP009444-RA | Su(var)205 | 186 |
| AGAP001093-RA | Kayak | 167 |
| AGAP006392-RA | CG4617 | 132 |
| AGAP005655-RA | Cycle | 130 |
| AGAP004990-RA | Mbf1 | 119 |
| AGAP006642-RA | Dve | 115 |
| AGAP006022-RA | Met | 110 |
| AGAP010405-RA | Maf-S | 107 |
| AGAP009676-RA | Chameau | 103 |

Enrichment analysis was run for every transcription factor and associated interactors for all GO term categories (Ashburner et al., 2000), KEGG pathways (Kanehisa and Goto, 2000), gene families previously associated with resistance (Balabanidou et al., 2016; Ingham et al., 2018, 2019; Müller et al., 2008; Voice et al., 2015) and Reactome pathways based on *Drosophila* homology (Jassal et al., 2020) (Table 1, Supplementary Table 1).

GO enrichments were present for 19/44 of the transcription factors across all ontology categories (Molecular Function, Cellular Component and Biological Process). A large number of GO terms were significant across different transcription factor interactions analysed; however, the terms were largely non-overlapping indicating that the transcription factors are playing differing roles in insecticide response (Supplementary Table 1, Supplementary Figure 2). Just three GO terms (membrane-enclosed lumen, organelle lumen, intracellular organelle lumen) were significant across four transcription factors and relate to localisation rather than function of the associated transcripts.

KEGG enrichments were present for 41/44 transcription factors (Supplementary Table 1, Supplementary Figure 3), again there was minimal overlap in the enriched pathways, in agreement with the divergent enriched GO terms. Three KEGG pathways were significant for five transcription factor associations (Valine, leucine and isoleucine degradation, Metabolism of xenobiotics by cytochrome P450 and Drug metabolism - cytochrome P450) and six terms were significant for four transcription factor associations (RNA polymerase, Peroxisome, Oxidative phosphorylation, Longevity regulating pathway - multiple species, Fatty acid degradation, Drug metabolism - other enzymes).

Given our *a priori* knowledge of insecticide resistance, enrichment analysis was also carried out for detoxification gene families, the cuticular hydrocarbon synthesis pathway and chemosensory proteins; three well described resistance mechanisms (Balabanidou et al., 2016; Ingham et al., 2019; Müller et al., 2008; Voice et al., 2015). Enrichments for these families occurred in 25/44 transcription factors with cytochrome p450s being significantly enriched in nine, GSTs in eight, UGTs in six, COEs in five ABCs in three, chemosensory proteins in three and the cuticular hydrocarbon pathway in five (Supplementary Table 1, Supplementary Figure 4). Reactome enrichment was also carried out, with significance for at least one pathway in 23/44 of the transcription factors (Supplementary Table 1, Supplementary Figure 5).

Taken together, these data indicate that the applied DBN is successfully capturing differing roles of the transcription factors in insecticide exposure response and the enrichment of a large number of *a priori* detoxification candidates indicates we are successfully capturing transcription factors controlling metabolic response to insecticide exposure.

### Key transcriptional regulators of insecticide response

Transcription factors that have previously been implicated in insecticide resistance or stress response and those that have interactors which show a clear functional enrichment from the above analysis are described in greater detail below.

#### Diminuitive

*Diminuitive* (*Dm*, AGAP00646-RA) is a central network hub with 702 interactors and its interactors are enriched in multiple KEGG pathways such as fatty acid degradation, nicotinate and nicotinamide metabolism and starch and sucrose metabolism (Supplementary Table 1). Within the interactome are two transcript variants of *para* the target site for pyrethroid insecticides (Martinez-Torres et al., 1998); AGAP004707-RA and AGAP004707-RC with posterior probabilities of 0.38 and 0.26 respectively. Eleven further transcription factors are within the *Dm* interactome including *cnc*, the transcriptional co-factor of *Maf-S*, previously linked with resistance (Ingham et al., 2017) and *run*, a transcription factor involved in neuronal development; both these interactions have been demonstrated in Drosophila (Kappes et al., 2011; Nagy et al., 2013). *Bap111* and *net*, two of the 14 transcription factor hubs identified above are also found within the *Dm* interactome. Previous work has demonstrated that attenuating *Dm* expression in *An. gambiae* results in significantly higher mortality post-pyrethroid exposure (Ingham et al., 2018); this role is underlined by significant enrichment of detoxification gene families in this cluster including cytochrome p450s (p = 0.011), carboxylesterases (p= 0.019) and GSTs (p = 1.35e-5) (including the GSTE cluster associated with pyrethroid resistance (Riveron et al., 2014; Wilding et al., 2014)). Additionally, the cytochrome p450 reductase, cpr, is represented in this cluster. *Dm* is also enriched in members of the cuticular hydrocarbon synthesis pathway (p = 3.2e-4) including *CYP4G16, CYP4G17* which catalyse the final step of CHC biosynthesis (Balabanidou et al., 2016). As part of the network validation, *Dm* expression was attenuated and shown to influence the expression of both *CYP4G16* (AGAP001076) and *CYP4G17* (AGAP000877), resulting in large reductions in their expression, further indicating a potential role for *Dm* in cuticular hydrocarbon synthesis.

#### *Met* and *Maf-S*

Both *Maf-S* (AGAP010405-RA) and *Met* (AGAP006022-RA) have previously been shown to have important roles in insecticide response (Ingham et al., 2018, 2017). In the absence of insecticide exposure, attenuation of expression of these transcripts demonstrated that both influenced the expression of key pyrethroid metabolisers such as *CYP6M2, CYP6Z2, CYP6Z3, CYP6P4, GSTD1* and *CYP9K1* (Yunta et al., 2019). In this analysis, *Met* is a direct target of *Maf-S*, in agreement with a previously published *Maf-S* knockdown array (Ingham et al., 2017). *Met* shows significant enrichment for cytochrome p450s (p = 2.5e-4), including *CYP6Z1, CYP6Z2* and *CYP6Z3* the latter two of which are amongst the most strongly induced p450s in the dataset. Interestingly, *Maf-S* does not show enrichment in *a priori* detoxification families, being enriched only in the KEGG pathway mitophagy (p = 0.037), indicating *Maf-S* may play a role in selective degradation of mitochondria following insecticide induced damage. Interestingly, the *Maf-S* pathway has been shown to be involved in mitophagy resulting from reactive oxygen species damage in mammalian systems through the action of *Pink1* (Murata et al., 2015), a role conserved in *Drosophila* (Cornelissen et al., 2018). Examination of the interactome network shows that the *Anopheles* homolog of *Pink1* (AGAP004315-RA) is within the *Maf-S* associations with a posterior probability of 0.61, indicating that this role may be conserved in Anophelines.

#### Mbf1

*Multiprotein bridging factor 1* (*mbf1*, AGAP004990) has 119 interactors and is strongly enriched for GO terms related to the ribosome (p = 0.00031) and translation (p = 4.2e-5) and is highly enriched in the KEGG ribosome (p = 4.4e-5). The role of *mbf1* in *Drosophila* involved translocation to the nucleus upon cellular stress, where it serves as a co-activator of stress response genes (Jindra et al., 2004); despite this role no enrichment for detoxification transcripts is seen in the predicted *mbf1* associations. However, 3 chaperone proteins (*CCT3, 4, 6*), an oxidation resistant protein (AGAP001751-RA) are present in this network. AGAP002667 has the highest posterior probability in the network (0.827) and encodes the homolog of *Drosophila Tctp* which is necessary for genomic stability under genotoxic stress (Hong and Choi, 2013).

#### Capicua

*Capicua* (*cic*, AGAP003117-RA) has 269 interactors and strong enrichment in GO terms related to cell morphogenesis involved in neuron differentiation (p = 0.0018), cation channel activity (p = 0.000584) and synapse (p = 0.018). KEGG pathway shows the phosphatidylinositol signalling system to be enriched (p = 0.00016), further exploration of this pathway shows these enrichments to be around calcium signalling. Three nicotinic acetylcholine receptor subunits (*α3* (AGAP000329), *β1* (AGAP000966) and *α6* (AGAP002152)) are included in this network out of 10 receptor subunits. *Rdl* (AGAP006028) is also included in this network, the target for dieldrin insecticides and postulated to be a secondary target for pyrethroids (Taylor-Wells et al., 2015). Several homologs of *Drosophila* proteins involved in neuronal development and behaviour are also included in these associations, including: *dunce* (AGAP000236) which is involved in behavioural plasticity (Zhong and Wu, 2004), *RhoGAPp190* (AGAP000870), *RhoGAP100F* (AGAP003944) and *Ank2* (AGAP002272) which regulate axon growth and stability (Koch et al., 2008; Ng and Luo, 2004), and two homologs of *Gbeta13F* (AGAP005911 and AGAP005913) which are involved in neuronal cellular division (Schaefer et al., 2001).

#### Sugarbabe

*Sugarbabe* (*sug*, AGAP006736-RA) has 64 interactors and is significantly enriched in GO terms related to fatty acid metabolic process (p = 1.01-5) and fatty acid elongase activity (p = 1.97E-5); it is similarly enriched in the KEGG pathway fatty acid metabolism (p = 2.66E-5). The hydrocarbon synthesis pathway is highly enriched (p = 2.43E-8), as with *Root* a direct interactor of *Sug*, including 4 fatty acid elongases (AGAP007264, AGAP003195, AGAP003197 and AGAP013094). *Sug* acts as a control for the insulin pathway in Drosophila and so is involved in both lipid and carbohydrate metabolism (Varghese et al., 2010); this is reflected in both the enrichments for fatty acid metabolic processes and the presence of insulin-related transcripts in its interactome, such as AGAP002544, the homolog of Drosophila *svp* which regulates insulin signalling (Musselman et al., 2018), the *AkhR* homolog (AGAP002156-RA) which encodes a G-protein coupled receptor that modulates lipid and carbohydrate metabolism (Bharucha et al., 2008) and *Lsd-1* related to lipolysis and lipid storage (Beller et al., 2010).

## Discussion

In this study, we apply a dynamic Bayesian network approach to whole transcriptome time-course data post-sublethal exposure of *An. coluzzii* to the pyrethroid insecticide deltamethrin (Ingham et al., 2020). The modified DBN model employed here allows correction for not only circadian rhythms, but also for mosquito ageing, a critical variable in the resistance status (Jones et al., 2012). Interactions predicted by this model were then validated *in vivo*, demonstrating high model confidence, with 70% precision. The high model precision and the overlapping biological functions with known transcription factors in *Drosophila* demonstrates the utility of this approach in assigning transcription factor function. Furthermore, this study highlights the potential for use of this methodology across multiple species of interest in which lower resolution time points are more feasible than those seen in model organism studies. Potential applications of this methodology could include exploring transcriptional regulation of pesticide response in other pest species or exposing the same species to additional stressors to distinguish between transcription factors involved in general and insecticide induced stress response.

In this study we highlight 44 transcription factors with putative roles in response to sublethal pesticide exposure, 41 of which have not previously been linked to insecticide resistance. Of the 8832 transcripts differential in the data set used, 4132 transcripts were assigned associations with these 44 transcription factors, using a posterior probability cut-off of >0.25. The assignment of 47% of the overall responsive transcripts is likely due to necessity of reducing the number of transcription factors to less than 50 transcripts and responsive transcripts being regulated by other mechanisms such as non-coding regulatory machinery. The transcription factors selected here for further analysis were identified by applying an SILGGM model (Zhang et al., 2018) to 28 insecticide resistant vs susceptible microarray datasets performed on the *Anopheles gambiae* species complex collated by Ingham et al. (Ingham et al., 2018) and exploring enrichments of co-correlated transcripts; this represents a confounding aspect of this methodology as these transcripts are constitutively overexpressed and not induced by insecticide exposure due to the nature of the transcriptomic designs.

Of the 44 transcription factors, 3 had previously been linked with insecticide resistance in *Anopheles* mosquitoes and just 11 had been previously studied in mosquito species in any context (Amenya et al., 2010; Chen et al., 2017; Chowdhury et al., 2020; Fu et al., 2020; Ingham et al., 2018, 2017; Luna et al., 2006; Maliti et al., 2016; Ruiz et al., 2019; Wang et al., 2017; Wülbeck and Simpson, 2002). All but 4 of these transcription factors have a well-defined role in *Drosophila*. Using a posterior probability cut off of >0.25, the number of associations showed high levels of variation with an average edge count of 107.84±150.42, potentially demonstrating differential importance in insecticide response, with those transcription factors with a high number of edges or high network connectivity being more important. Fourteen transcription factors were designated as transcript ‘hubs’ based on high levels of network interconnectivity (>100 edges), a further 11 transcription factors had over 50 associations.

Enrichment analysis was performed for all transcription factors in the network, using GO Terms, KEGG Pathway, Reactome and *a priori* transcript families with links to resistance. Interestingly, the overlap of enriched terms was low, indicating that each transcription factor may play a differing role in the response to insecticides. 25 transcription factors show enrichments in *a priori* gene families; this may be an unsurprising feature of this dataset given the obvious change in expression across multiple members of these families within this dataset and their documented importance in insecticide metabolism (Ingham et al., 2020). GO terms enriched across multiple transcription factors include terms expected in an insecticide response, such as xenobiotic response by cytochrome p450s, drug metabolism, peroxisome and longevity. The former two enrichment terms are in agreement with the well-established dogma that up-regulation of members of the cytochrome p450 class play a direct role in increasing the rate of insecticide metabolism (Ingham et al., 2018; Yunta et al., 2019). Both peroxisome and longevity are involved in response to reactive oxygen species, which is likely to be an *in vivo* response to pyrethroid exposure as shown indirectly in mosquitoes (Oliver and Brooke, 2016) and directly in mammalian cells (Wang et al., 2016). Interestingly, changes to the respiratory pathway through alterations to the oxidative phosphorylation pathway also appears across multiple transcription factors and is a striking feature of this dataset (Ingham et al., 2020).

To cross-validate the function of these transcription factors, their known functions in the model organism *Drosophila* were explored. Despite the differences in hypotheses explored in this study and the available data in discerning *Drosophila* pathways, there were clear overlaps in transcription factor roles and associations. For example, *Dm* is known to play a role in lipid and glucose homeostasis in *Drosophila* (Parisi et al., 2013) and here, the associations are enriched in the KEGG pathways starch and sucrose metabolism and fatty acid degradation; this is similar to *Hnf4* which is involved in lipid mobilisation (Palanker et al., 2009) and is enriched in the GO term related to lipid transporter activity. Several further transcription factors show overlap with *Drosophila* function, including *Sugarbabe* which is involved with the insulin pathway and regulation of lipids (Varghese et al., 2010), *dve* which is involved in mitochondrial reactive oxygen species modulation (Baqri et al., 2014), *Ets21C* which is a stress-inducible transcription factor (Mundorf et al., 2019), *klumpfuss* whose role is related to cell death and the mitochondria and *Maf-S* and its role in mitophagy (Murata et al., 2015). Interestingly, *Hnf4* has had its role in lipid metabolism confirmed in mosquito studies (Wang et al., 2017) and *Pangolin* has been shown to be involved in chromatin remodelling upon *Plasmodium* infection (Ruiz et al., 2019), mirroring the enrichment seen here for chromatin and DNA-packing GO term enrichments, adding further confidence to the predicted associations.

This study provides not only previously unreported transcription factors that are involved in the transcriptional response to pesticide exposure but demonstrates the utility of applying a model-based approach to lower-resolution time course data in ascertaining these associations. Here, six transcription factors and their interactomes were delineated as hub transcripts within the network, all of which have either been previously linked to resistance or stress response in *Anopheles* (*Dm, Maf-S* and *Met*) (Ingham et al., 2018, 2017) or *Drosophila (mbf1*) (Jindra et al., 2004) or are highly significantly enriched for clear functions (*capicua* and *sug)*. These transcription factors are likely to be involved in different facets of insecticide response and represent pathways that should be further explored. The modelling approach taken here, which accounts for both circadian patterns and ageing, two key determinants in pesticide resistance, can be applied widely to other pest or vector species. Using this approach will provide invaluable information on changes to pest biology post-pesticide exposure and will elucidate new pathways to characterise and target to tackle the ongoing threat of pesticide resistance.

## Materials and Methods

### Microarray Experiments

Microarrays were taken from (Ingham et al., 2020) and consist of deltamethrin exposed mosquitoes compared to unexposed at the following time points post-exposure: 0 minutes, 30 minutes, 1 hour, 2 hours, 4 hours, 8 hours, 12 hours, 24 hours, 48 hours and 72 hours. To account for ageing effects, a twin time course was performed using age matched females that were unexposed to insecticide at the following time points: 8 hours, 12 hours, 24 hours, 48 hours and 72 hours. All mosquitoes within one experimental time course came from the same generation. Experimental data is available on exposure time course (E-MTAB-9422) and ageing time course (E-MTAB-9423).

### Transcription factor identification

28 microarray datasets encompassing resistant vs susceptible members of the *Anopheles gambiae* species complex were used from Ingham et al. 2018 (Ingham et al., 2018). A de-sparsified node-wise scaled lasso (Janková and van de Geer, 2018, 2017) implemented in the R package SILGGM (Zhang et al., 2018), was used to infer the gene network. This method employs L1-regularisation to preserve sparsity in the estimated network. For the L1-regularisation, the default value of the tuning parameter λ, was used: √log(p)/n, where p is the number of variables and n is the number of samples. The resultant Gaussian graphical model produced a 14079×14079 file for every possible interaction in the transcriptome. Each interaction had an associated p-value for precision (Supplementary Table 4). A cut-off value of p ≤ 0.1 was used to filter all interactions to prevent loss of potentially interesting transcription factors due to the differing experimental design of the data set used. Annotated Drosophila transcription factors were downloaded from FlyTF (Adryan and Teichmann, 2006) (http://flytf.gen.cam.ac.uk/) and *Anopheles* homologs identified using FlyMine (Lyne et al., 2007) (https://www.flymine.org) resulting in 559 putative transcription factors; all 559 were then extracted from the inferred network with all associated putative co-correlating transcripts. clusterProfiler (Yu et al., 2012) and AnnotationForge (Carlson and Pagès, n.d.) were used to perform GO enrichments using an Anopheles database built from PEST/VectorBase (Giraldo-Calderón et al., 2014) on Biological processes on transcription factors with > 10 interactors. Transcription factors enriched in the following character patterns were extracted: ‘stress’; ‘oxi’; ‘lipid’; ‘behaviour’; ‘response’; ‘fat’; ‘sensory’ and ‘ATP’ leading to 54 transcription factors. The transcription factors were further filtered on at least 50% of the transcripts in the cluster generated by SILGGM being differentially expressed in at least 1 time point within the time course datasets. This procedure resulted in 44 transcription factors being retained.

### Network reconstruction using Dynamic Bayesian Networks

Dynamic Bayesian networks (DBNs) (Dondelinger et al., 2013) were used to identify directed associations between the transcription factors and putative regulated genes. A dynamic Bayesian network defines a graphical model for the dynamics of time series data, where the gene expression *X*_*i*_(*t*) of gene i at time t depends on the gene expression *X*_*j*_(*t*) of all transcription factor genes j at time t − δ. The relationship can be described by the following auto-regressive linear regression:

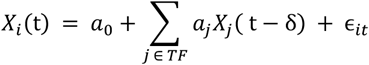

where 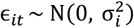, and TF is the set of indices representing the transcription factors. To impose regularisation, we assumed truncated Poisson priors on the number of regression parameters *a*_*j*_ that are non-zero:

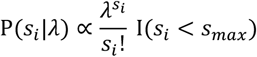

where *s*_*max*_ is the maximum number of transcription factors regulating a single gene. We set *s*_*max*_ = 5. Conditional on *s*_*i*_, the number of non-zero transcription factor-gene associations, the prior on the set of transcription factors for a given gene is simply a uniform distribution.

Inference of the network structure can be done via a Markov Chain Monte Carlo algorithm. For full details on the model and inference procedure, please see (Dondelinger et al., 2013). Note that here we employ a simplified version of this model which does not use a changepoint model or information sharing priors.

Prior to applying the network inference model, we preprocessed the log-fold change data by first averaging the values for genes with multiple probes to obtain one measurement per gene. We then employ LOESS estimation to interpolate the time points at t − δ, where we choose δ = 0.5 hours as the time interval. Interpolation is necessary, as the DBN method requires equal time intervals between each pair of measurements to estimate consistent associations.

We further extend the model to correct for circadian rhythms and ageing effects in the gene expression levels. For the circadian rhythm correction, we assume that all circadian rhythms have a period of 24 hours, and augment the design matrix **X** = {*X*_1_(*t*), …, *X*_*p*_(*t*)} with two additional columns for the sine and cosine functions of a 24-hour periodic signal:

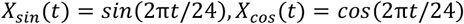

The resulting harmonic regression model with automatically correct for circadian rhythms by including the periodic signal as a parent in the network, while non-periodic genes will remain unconnected to this signal.

Similarly, we add additional columns for the data arising from the ageing controls to correct for the effect of ageing. Note that here we only have data starting from 8 hours, so earlier time points will be uncorrected, and the corresponding values in the design matrix will be set to zero. The final autoregressive model looks as follows:

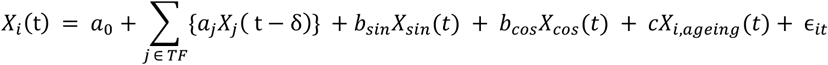

where *X*_*i,ageing*_(*t*) is the log-fold change of the ageing controls.

We summarize the results of the DBN analysis using the marginal posterior probability of each transcription factor – target gene association, which can be calculated by obtaining samples from the converged Markov chain and averaging over the presence/absence status of each edge. In order to determine a sensible threshold for the marginal posterior probability that keeps the false discovery rate low, we implement the following permutation test to estimate the posterior probabilities under the null hypothesis of no associations: for each of n=500 iterations, we randomly permute the log-fold changes for one transcription factor. Any associations with the target gene should then be entirely by chance. Taking all n=500 samples of the null distribution obtained in this way, we determine that a threshold of 0.23 was sufficient to only produce one false positive out of 500 randomizations, or a false positive rate of 0.002. The network was displayed using Cytoscape (Shannon et al., 2003).

### NetworkVis App

The NetworkVis app and associated data can be downloaded on Github (https://github.com/VictoriaIngham/NetworkVis_TimeCourse) and installed as described. ShinyR (Chang et al., 2017) was used to create a user interface, both VisNetwork (Almende et al., 2018) and igraph (Csardi and Nepusz, 2006) were used to allow dynamic selection of nodes and edges, and to display the network.

### Enrichment analysis

Enrichment analysis was performed using clusterProfiler (Yu et al., 2012) and a custom *Anopheles* database produced using AnnotationForge (Carlson and Pagès, n.d.). GO term and KEGG enrichments were performed using a Benjamini-Hochberg corrected p value cut-off of ≤ 0.05 with transcription factors > 10 interactions. Clusters of each transcription factor were compared using the compareCluster function; these were then displayed using Cytoscape (Shannon et al., 2003). Enrichment analysis on individual gene families were performed using a hypergeometric test with the phyper function in R; significance was considered when p ≤ 0.05. Reactome analysis was also performed using a hypergeometric test with p ≤ 0.05; *Drosophila* pathway membership was downloaded from Reactome.org (https://reactome.org/) (Jassal et al., 2020) for each pathway of interest, FlyMine (Lyne et al., 2007) was then used to convert these to *Anopheles* homologs. FlyBase (Consortium, 2003) was used to determine functions of homologs throughout the analysis.

### Validation of Network

We first performed a simulation study to determine the number of associations that need to be tested experimentally in order to obtain an accurate estimate of the precision of our network inference method. We made the following assumptions: (i) The mean number of gene regulated by each transcription factor is 10, and the actual number of regulated genes follows a Poisson distribution; (ii) The rate of true positives (correctly predicted associations) of our network is 0.75, and the rate of true negatives (correctly predicted non-associations) is 0.997; this results in a precision of ∼0.6 and a recall of ∼0.72; (iii) Transcription factors and regulated genes to test are selected randomly and (iv) The qPCR knockdown test is 100% accurate. The results of the simulation study can be found in Appendix 1. We concluded that testing 4 regulatory relationships for 7 transcription factors has a 70% chance of obtaining an estimate of the precision that falls within 10% of the true precision, and a 95% chance of obtaining an estimate that falls within 20% of the true precision.

In order to choose associations for validation, we then chose interactors by extracting the transcription factor of interest and associated transcripts from the results of the network inference. Transcripts were listed as 1 to n based on posterior probability in descending order. A random number generator was then used to select 4 transcripts for validation from 7 transcription factors chosen based on previous knockdown in the case of *Maf-S, Met, Dm* or through a random number geenerator.

### Mosquito Rearing

The *An. coluzzii* VK7 colony reared and profiled at Liverpool School of Tropical Medicine were used for all experiments (Williams et al., 2019). VK7 are a highly pyrethroid resistant population originating from Vallée de Kou, Burkina Faso (Toé et al., 2015). They have been reared at LSTM since 2014 under pyrethroid selection pressure (Williams et al., 2019). All mosquitoes used were reared under standard insectary conditions of 27oC and 70-80% relative humidity under a 12:12 photoperiod and are presumed mated.

### dsRNA knockdown

RNAi was performed using 7 transcription factors based on previous publication of knockdown (Maf-s, Met, Dm (Ingham et al., 2018, 2017)) or through random selection using a random number generator (Med, Pan, l(1)sc, mbf1) (Supplementary Table 4). PCR was performed on 3-day old VK7 unexposed cDNA using Phusion^®^ High-Fidelity DNA Polymerase (Thermo Scientific) following manufacturer’s instructions and primer sets with a T7 docking sequence at the 5′ end of both the sense and antisense primers (Supplementary Table 4). Primers were designed as previously described (Ingham et al., 2018). PCR was performed using the following cycles: 98oC for 30s, (98oC 7s, 65oC 10s, 72oC 10s) x35 and 72oC 5 minutes. PCR product was then purified using a Qiagen QIAquick PCR Purification Kit following manufacturers’ instructions. dsRNA was then synthesised using a Megascript^®^ T7 Transcription (Ambion) kit, following manufacturer’s instructions (16-hour 37 °C incubation). The dsRNA was cleaned using a MegaClear^®^ Transcription Clear Up (Ambion) kit, with DEPC water, twice heated at 65 °C for 10 min, to elute the sample. The resultant dsRNA product was analysed using a nanodrop spectrometer (Nanodrop Technologies, UK) and subsequently concentrated to 3 μg/μl using a vacuum centrifuge at 35oC. 69nL of dsRNA was subsequently injected into presumed mated, non-blood fed, 3-day old VK7 females immobilised using a CO_2_ block using a NanoInject II. 50 females were injected with each of the transcription factor dsRNA and 50 with dsGFP as a non-endogenous control.

### Insecticide Exposures

25-30 female mosquitoes were exposed to 0.05% deltamethrin impregnated papers for one hour in a standard tube bioassay kit following WHO guidelines. Post-exposure mosquitoes were transferred into holding tubes and maintained on sucrose solution.

### RNA extraction and cDNA synthesis

RNA was extracted from 7-10 female mosquitoes in biological triplicate for each experimental group. RNA was extracted from homogenised mosquitoes using a PicoPure RNA isolation kit (Thermo Fisher, UK) following manufacturers’ instructions and treated with DNAase (Qiagen) to remove any DNA contamination. Quality of RNA was checked using a nanodrop spectrophotometer (Nanodrop Technologies UK). 1-4µg of RNA from each experimental set was reversed transcribed using OligoDTT (Invitrogen) and Superscript III (Invitrogen) according to manufacturers’ instructions. The following experimental groups were used: (i) knockdown efficacy for each transcription factor and the GFP control using females 48-hours post RNAi injection and (ii) pathway validation using females 48-hours after they were exposed to 0.05% deltamethrin for 48-hours post-injection for transcription factors and GFP controls.

### qPCR validation

Quantitative real-time PCR was performed using SYBR Green Supermix III (Applied Biosystems, UK) using an MX3005 and the associated MxPro software v4.10 (Agilent, UK). Primer Blast (NCBI) was used to design primer pairs. Where possible, primers were designed to span an exon junction (Supplementary Table 4). Each 20µl reaction contained 10µl SYBR Green Supermix, 0.3µM of each primer and 1µl of 4ng/µL cDNA. Standard curves for each primer set were used to calculate efficiency, using five 1:5 dilutions of cDNA to ensure that all primer sets met the MIQE guidelines (90-120% efficiency) (Bustin et al., 2009). qPCR was performed with the following conditions: 3 minutes at 95oC, with 40 cycles of 10 seconds at 95oC and 10 seconds at 60oC. Relative expression was normalised against two housekeeping genes: EF (AGAP005128) and S7 (AGAP010592) and analysed using comparative CT method (Schmittgen and Livak, 2008). qPCR was used to determine the efficacy of transcription factor knockdown by comparing cDNA taken from mosquitoes 48-hours post dsRNA injection for each transcription factor and comparing it to GFP-injected controls all taken from the same mosquito generation. To validate findings in the network, qPCR was performed on dsRNA injected mosquitoes exposed to 0.05% deltamethrin at 48-hours post injection, these mosquitoes were then left for a further 48-hours before harvesting; again, transcription factor injected mosquitoes were compared to the dsGFP injected controls.

## Code Availability

Code used for analysis in this study is available on Github. Network visualisation is available at https://github.com/VictoriaIngham/NetworkVis_TimeCourse, model code is available on the CRAN repository: https://cran.r-project.org/web/packages/EDISON and full analysis is available at https://github.com/FrankD/AnophelesInsecticideExposure.

## Acknowledgements

This work was supported by a Medical Research Council Skills Development Fellowship (MR/R024839/1) to VAI. We thank Hilary Ranson and David Weetman for valuable feedback on the manuscript and Jessica Carson, Marion Morris and Ruth Cowlishaw for insectary support.

## Author Contributions

VAI and FD designed and implemented the experiment. SCN performed the SILGGM analysis, FD modified and implemented the dynamic Bayesian network, SE provided rearing, bioassay and molecular biology support, VAI performed the lab-based experiments and analysed all data. VAI and FD drafted the manuscript.

## Competing Interests

The authors declare that they have no competing interests.

## Data Availability

The datasets used in this experiment are available at ArrayExpress under E-MTAB-9422 and E-MTAB-9423. The authors declare that all other data supporting the findings of this study, are available within the article and its Supplementary Information files or are available from the authors upon request.

## Notes

### Competing Interest Statement

The authors have declared no competing interest.

