## Supplementary figures and images for "Capturing the transcription factor interactome in response to sub-lethal insecticide exposure"

### Supplementary Figure 1

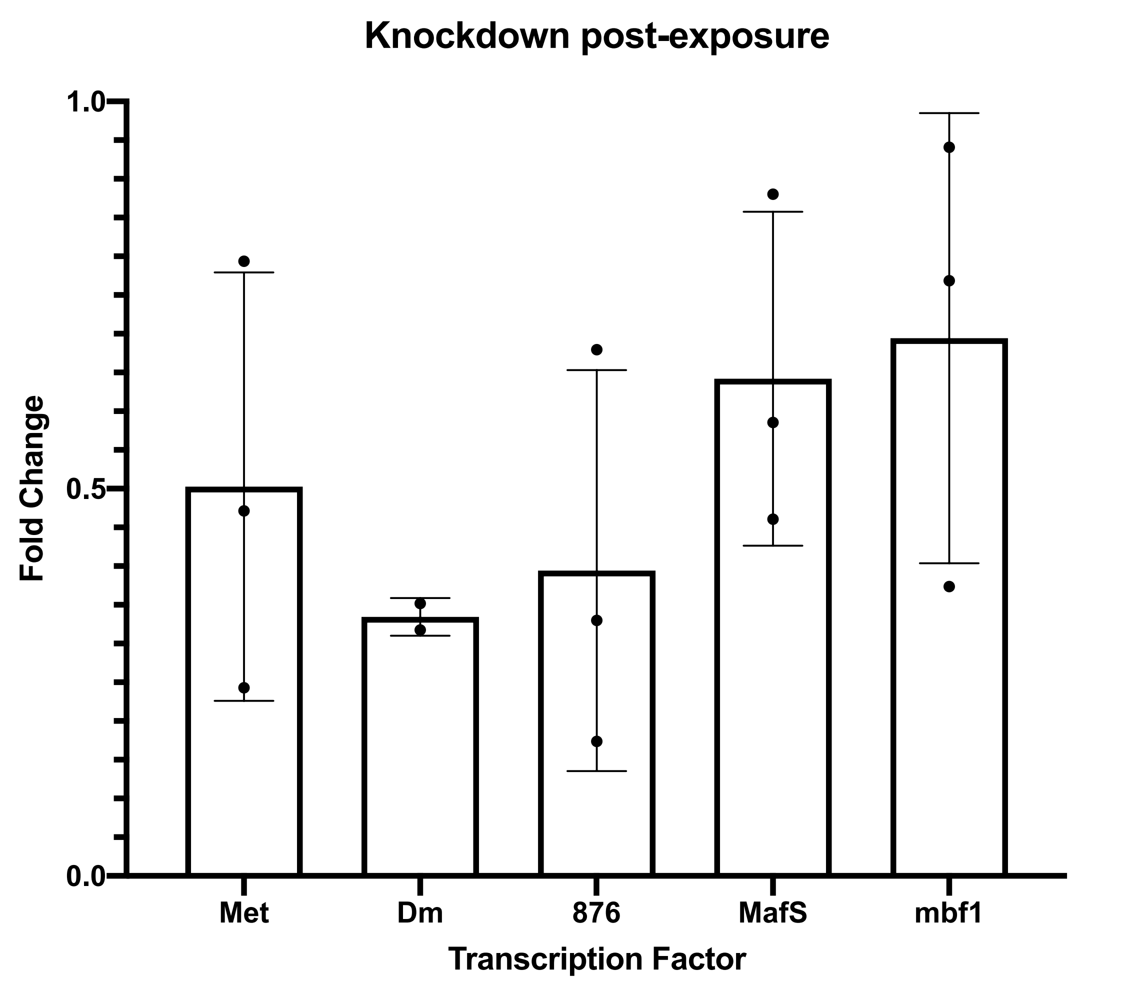

### Supplementary Figure 2

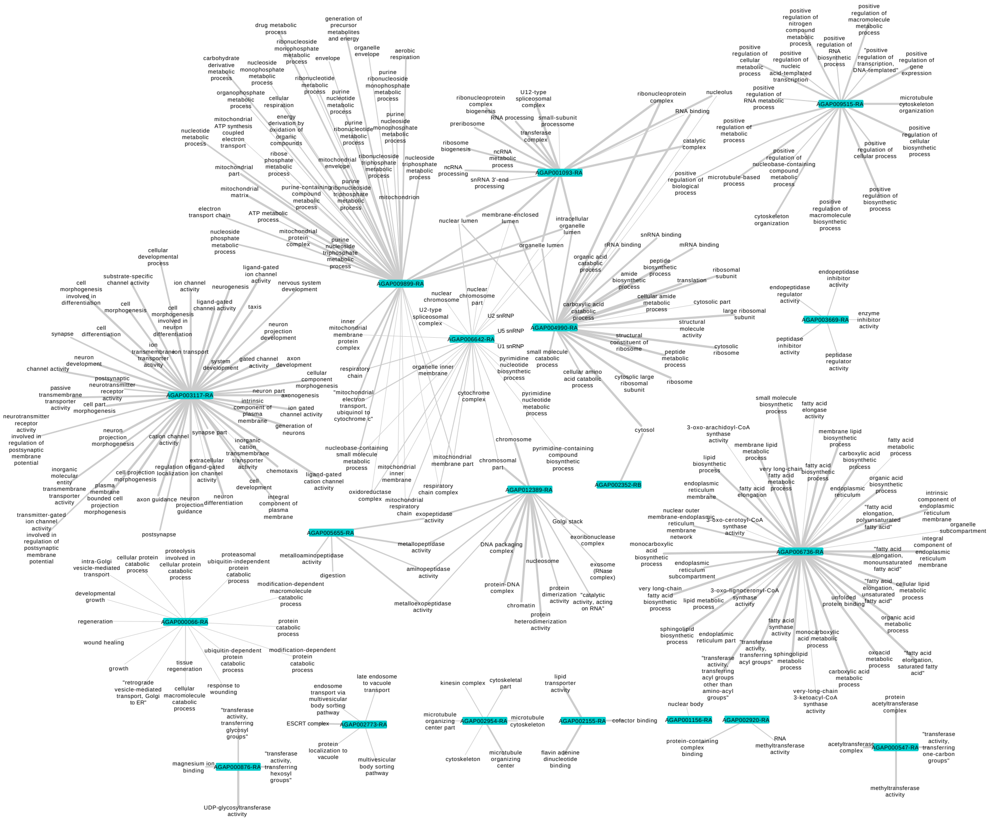

### Supplementary Figure 3

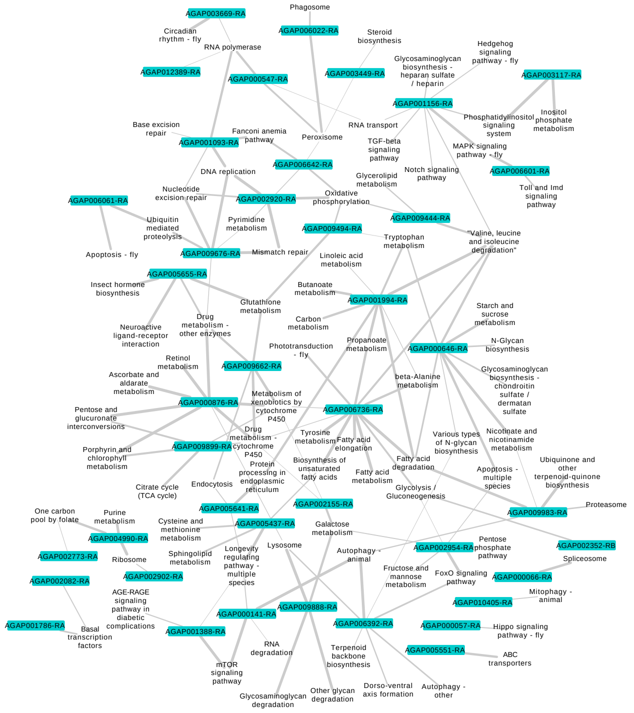

### Supplementary Figure 4

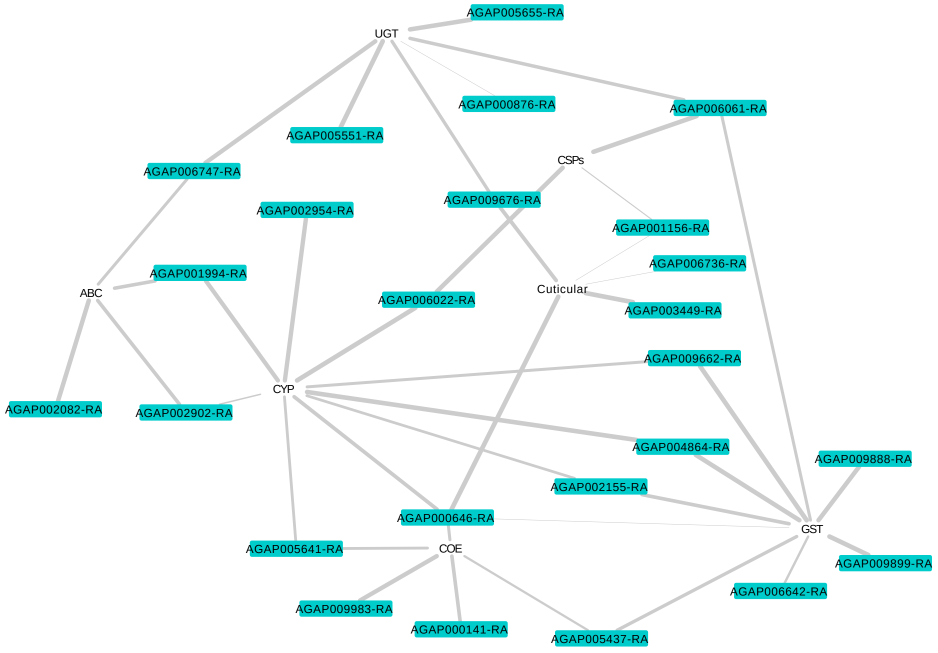

### Supplementary Figure 5

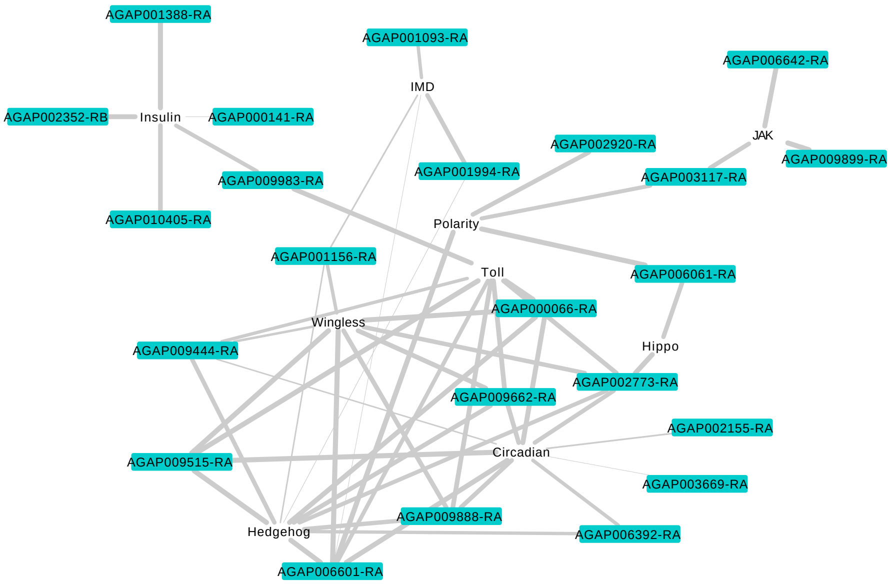
