## Supplementary Figure Legends for "Capturing the transcription factor interactome in response to sub-lethal insecticide exposure"

**Supplementary Table 3: Primer List.** dsRNA and qPCR primers used in the study, names and sequences. _F refers to the forward primer and _R the reverse

**Supplementary Figure 1: Transcription factor knockdown.** qPCR of relative fold change of transcription factor dsRNA injections compared to dsGFP injected control 48 hours post-exposure.

**Supplementary Figure 2: Visualisation of GO term enrichments.** Significant (p < 0.05) GO terms for each transcription factor (blue). Thickness of the edge represents significance, with thicker edges having smaller p values.

**Supplementary Figure 3: Visualisation of KEGG term enrichments.** Significant (p < 0.05) KEGG terms for each transcription factor (blue). Thickness of the edge represents significance, with thicker edges having smaller p values.

**Supplementary Figure 4: Visualisation of detoxification families.** Significant (p < 0.05) detoxification families for each transcription factor (blue). Thickness of the edge represents significance, with thicker edges having smaller p values.

**Supplementary Figure 5: Visualisation of Reactome enrichments.** Significant (p < 0.05) Reactome terms for each transcription factor (blue). Thickness of the edge represents significance, with thicker edges having smaller p values.
